# ethoscopy & ethoscope-lab: a framework for behavioural analysis to lower entrance barrier and aid reproducibility

**DOI:** 10.1101/2022.11.28.517675

**Authors:** Laurence Blackhurst, Giorgio F. Gilestro

## Abstract

**Summary:** High-throughput analysis of behaviour is a pivotal instrument in modern neuroscience, allowing researchers to combine modern genetics breakthrough to unbiased, objective, reproducible experimental approaches. To this extent, we recently created an open-source hardware platform (ethoscope (Geissmann *et al*., 2017)) that allows for inexpensive, accessible, high-throughput analysis of behaviour in *Drosophila* or other animal models. Here we equip ethoscopes with a Python framework for data analysis, ethoscopy, designed to be a user-friendly yet powerful platform, meeting the requirements of researchers with limited coding expertise as well as experienced data scientists. Ethoscopy is best consumed in a prebaked Jupyter-based docker container, ethoscope-lab, to improve accessibility and to encourage the use of notebooks as a natural platform to share post-publication data analysis.

**Availability and implementation:** Ethoscopy is a Python package available on GitHub and PyPi. Ethoscope-lab is a docker container available on DockerHub. A landing page aggregating all the code and documentation is available at https://lab.gilest.ro/ethoscopy.

## Introduction

The use of open-source software in science is commended by researchers and funders alike, for it frees academic science from ties with possibly ephemeral third-parties; it creates a bottom-up environment in which researchers actively collaborate to build their own solutions based on their own demands; it maximises the value of public funding; it provides the transparency that is essential for reproducibility. At the same time, however, academic open-source software still encounters barriers in terms of ease of use, compatibility with multiple operating systems and constantly evolving software ecosystems, as well as often unmet good practices in post-publication data sharing. We recently created an open-source, well documented platform for high-throughput behavioural analysis in *Drosophila* that has been widely adopted by the community (ethoscope (Geissmann *et al*., 2017), https://lab.gilest.ro/ethoscope) and we initially equipped it with a powerful and versatile selection of R packages (rethomics (Geissmann *et al*., 2019)). However, while R remains a language of first-choice for data scientists, its structure, syntax and philosophy can prove unwelcoming to newcomers and ultimately discourage adoption in the field. Moreover, data science is constantly increasing its ties with machine learning and Python is the environment of choice to combine the two. Here we describe ethoscopy and ethoscope-lab, two instruments whose genesis was motivated by our very experience in the past years. These tools are not simply the porting of rethomics into a more accessible programming language: in fact, they offer new functions and tools, integrating with the modern sleep and circadian literature and at the same time adopt the best practice in code accessibility and data sharing. Here we detail how.

### Improved accessibility for the community at large

Software accessibility is a key aspect of bioinformatics, as even the best tools are useless if they are not adopted by the community. One way to improve accessibility is for authors to distribute Docker containers (Boettiger, 2015; Nüst *et al*., 2020). Docker containers increase adoption because they guarantee the tool will always ship with the needed dependencies; they are truly multi-platform as they offer the same experience to any user irrespective of the operating system they prefer; they guarantee that all researchers will operate on the same version of code and underlying libraries, ultimately increasing reproducibility. Ethoscope-lab is a docker container distributed via the official DockerHub platform and providing at any time access to the latest versions of ethoscopy and rethomics in a Jupyter instance.

### Improved accessibility for users within the laboratory

Ethoscope-lab ships with a stable release of JupyterHub, the multiuser web-based notebook that rapidly became the tool of choice for most data scientists (Perkel, 2018). The typical scenario will be to deploy ethoscope-lab on a shared powerful workstation, accessible to all laboratory members via the intranet or the internet. Ideally, the same workstation will also store the raw data of the behavioural analysis, as extracted directly by the ethoscopes, so that ethoscopy can have local, read-only access to them. Users will connect to Jupyter-hub using their favourite web-browser on a computer or tablet, login using their own credentials, and work in the cloud, saving their notebooks, stats and figures on a remote folder on the workstation that can be backup-ed up and shared backed with themselves or other users (for instance, via Samba or the open source software ownCloud). This scenario offers multiple advantages: all the users in the laboratory can access ethoscopy as a SaaS (Software as a Service) which is the best way to improve accessibility to end users; the docker container and the workstation can be setup and maintained by one tech-savvy user or by the IT department, allowing all other end users to concentrate on the data analysis, not the tool maintenance; the only machine requiring powerful computing specifications will be the workstation hosting the docker container so that users will be able to perform big-data analysis from any device and any location, saving time and resources.

### Good practice in post-publication data sharing

At the time of publication, researchers will share information on which version of the docker container they used for their analysis and upload fully annotated Jupyter notebooks as supplementary data. We provide several example notebooks that can serve as step-by-step tutorials on the most common functions of ethoscopy (Supplementary Material) and, at the same time, offer direct evidence of why a Jupyter notebook is considered the best instrument to share post-publication data processing in research. A reader that has access to: 1) the raw behavioural data, 2) the metadata describing the experimental conditions, 3) a well documented Jupyter notebook and 4) the matching version of ethoscopelab, will be able to reproduce and re-analyse and figure of any publication, with no risk of obsolescence even after decades. All these can be shared on the journal platform or via third party scholarly services, such as Zenodo.

### Ethoscopy structure and features

Similarly to rethomics, ethoscopy will work out-of-the-box on ethoscope and DAMs data (Rosato and Kyriacou, 2006) but can in principle process any behavioural data as long as they provide the three necessary descriptors of any behaviour: an animal identification (1) and at least one quantification of a behavioural trait (2) over time (3). Two common examples of tabular inputs are: a comma separated list associating an animal identification (id column) to its Cartesian coordinates in space (x, y) over time (t); a spreadsheet associating animals id (id) to amount of food consumed per minute (f). As a proof of principle, we show how to import human actigraphy data collected through a commercially available device (FitBit Charge – Notebook 7 in the Supplementary Material). Once data are imported, the main structure in ethoscopy will build on a *pandas*.*DataFrame* (McKinney and others, 2010), the current handling standard in Python-based data science. A *behavpy* object combines two tabular structures, linking the raw behavioural data with their metadata and provides builtin custom methods and properties for handling data and plotting figures (Figure 1a). Because *behavpy* classes inherit directly from *pandas*.*DataFrame*, they also carry all upstream properties and methods of the DataFrame, thus providing access to the entire pandas toolbox. Inbuilt plotting wrappers can output figures either using the graphing library plotly (Sievert, 2020) or Seaborn (Waskom, 2021), a Matplotlib (Hunter, 2007) front end. These two open-source widely adopted plotting libraries come each with different strengths and weaknesses (Figure 1b). Plotly output is interactive by default and therefore particularly useful during exploratory data analysis: with a click of the mouse, users can zoom on smaller or larger windows of data or pick individual datasets, for instance hiding genotypes from a complex comparison to simplify the visualisation. Seaborn output, on the other hand, is static but lightweight and more easily customizable, and thus better suited for final data publication and sharing. In general, ethoscopy functions are mostly geared towards animals’ activity data collected with ethoscopes, but they are not limited to sleep. Here, for instance, we provide tutorials for quantification of sleep (Notebook 1 in Supplementary Material and Figure 2a-c), for circadian analysis (Notebook 3 in Supplementary Material and Figure 3a,b), for an analysis of sleep-depth that uses a hidden Markov model to identify sleep stages in *Drosophila* (Wiggin *et al*., 2020) (Notebook 2 in Supplementary Materials and Figure 1d,e) and, finally, for integration with highly comparative time-series analysis (HCTSA -(Fulcher and Jones, 2017)) either in its entirety (Notebook 5 in Supplementary Material) or its curated subset Catch22 (Lubba *et al*., 2019) (Notebook 6 in Supplementary Materials and Figure 3c,d).

**Figure 1.**
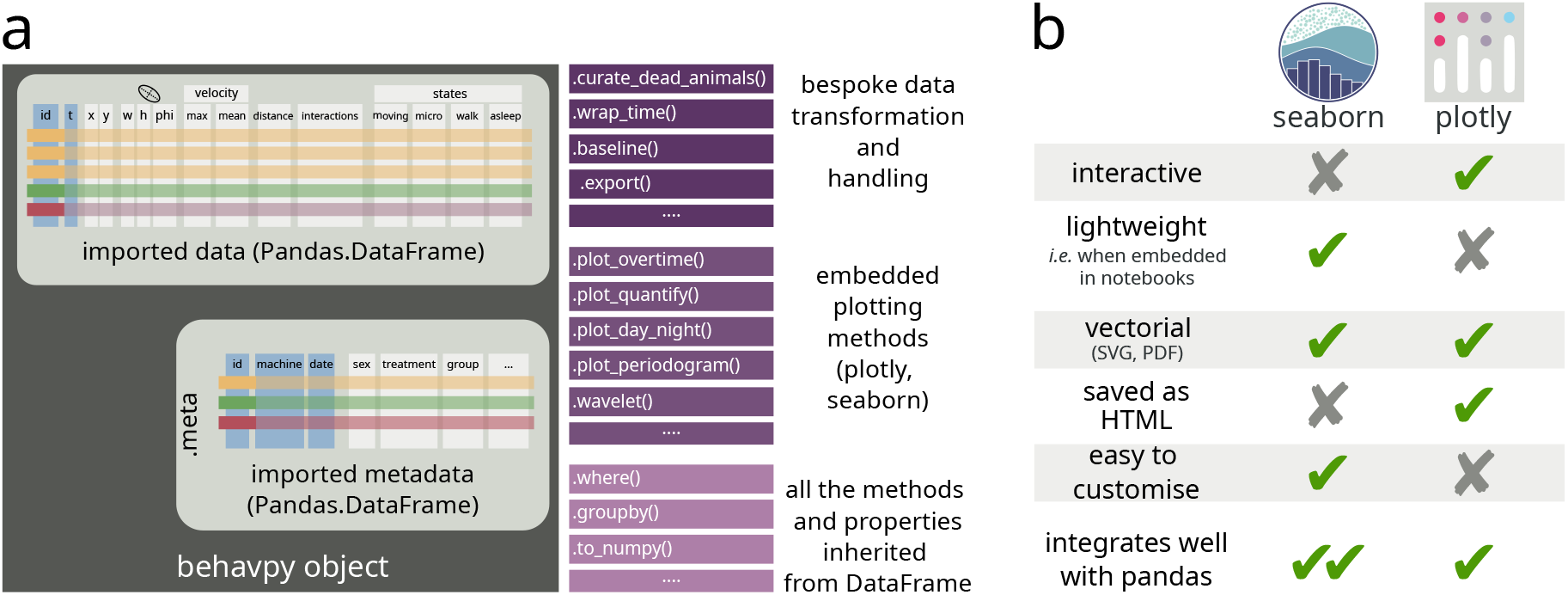
Ethoscopy structure and flexibility. **a)** Schematic structure a typical *behavpy* object. All *behavpy* classes are children of a *pandas*.*DataFrame* object and carry the behavioural time series as their main content. The main property of the class is a second *pandas*.*DataFrame*, .meta, with the associated metadata linked to the behavioural data through the *id* column. Any *behavpy* object features bespoke methods for data transformation or handling, builtin methods for plotting figures, as well as all the upstream methods and properties inherited from *pandas*.*DataFrame*. **b)** Panoptic summary of the strengths and weaknesses of the two default plotting libraries, seaborn and plotly.

**Figure 2.**
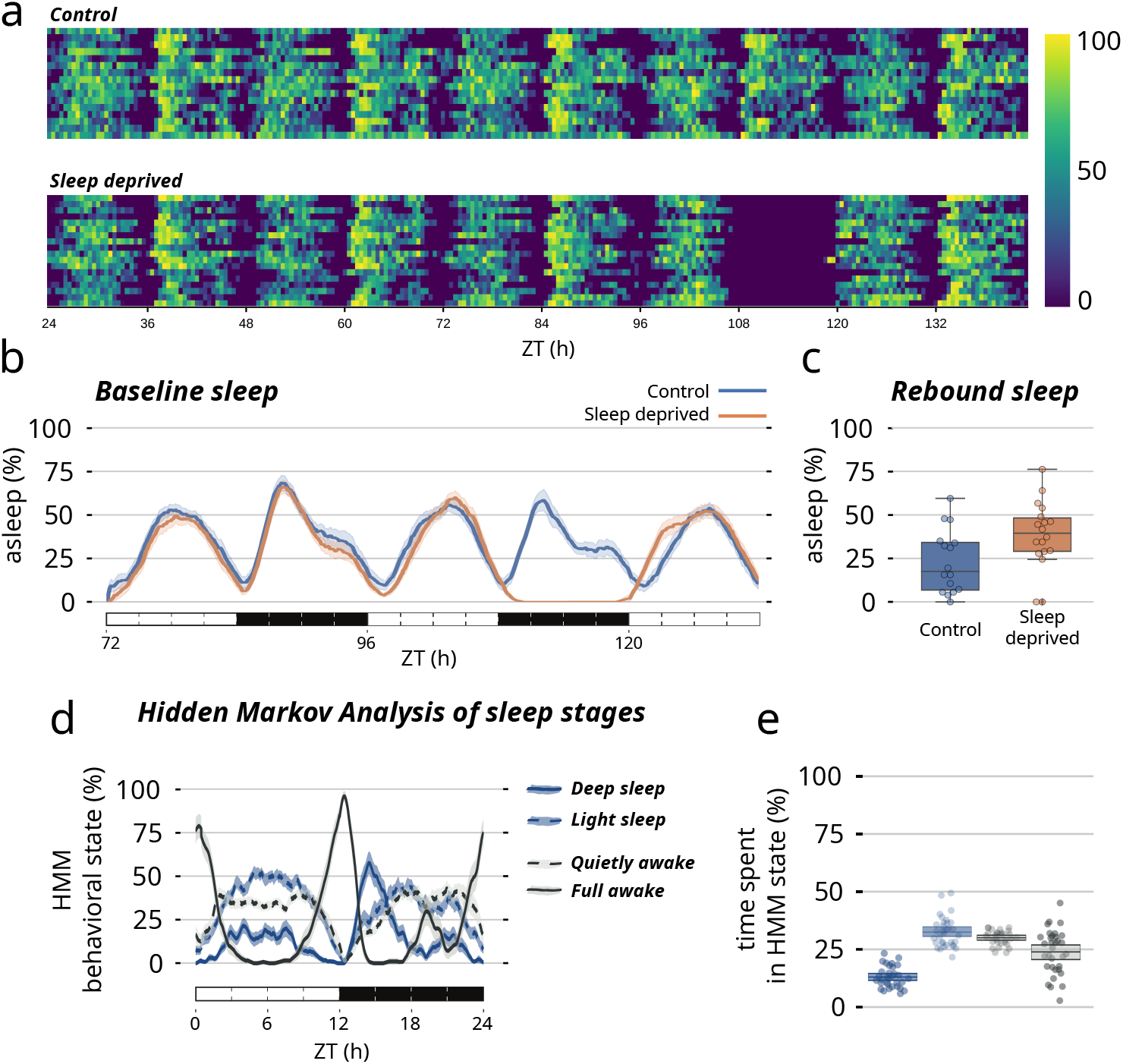
Ethoscopy examples of sleep analysis. **a)** Overview of sleep and activity using the heatmap plotting method of ethoscopy. Overall activity of a group of wild type flies in control conditions (upper panel) or during a sleep deprivation experiment (lower panel). The blue to yellow colour gradient quantifies activity (blue) and sleep (yellow). **b)** Sleep profile of the two experimental conditions over 2.5 days, including the sleep deprivation day. **c)** Quantification of rebound sleep for the experiment shown in b). **c)** Hidden Markov chains modelling of four covert sleep stages over the 24 hours period in a sample group of wild-type CantonS flies. ZT: Zeitgeber. **d)** quantification of each state over the 24h.

**Figure 3.**
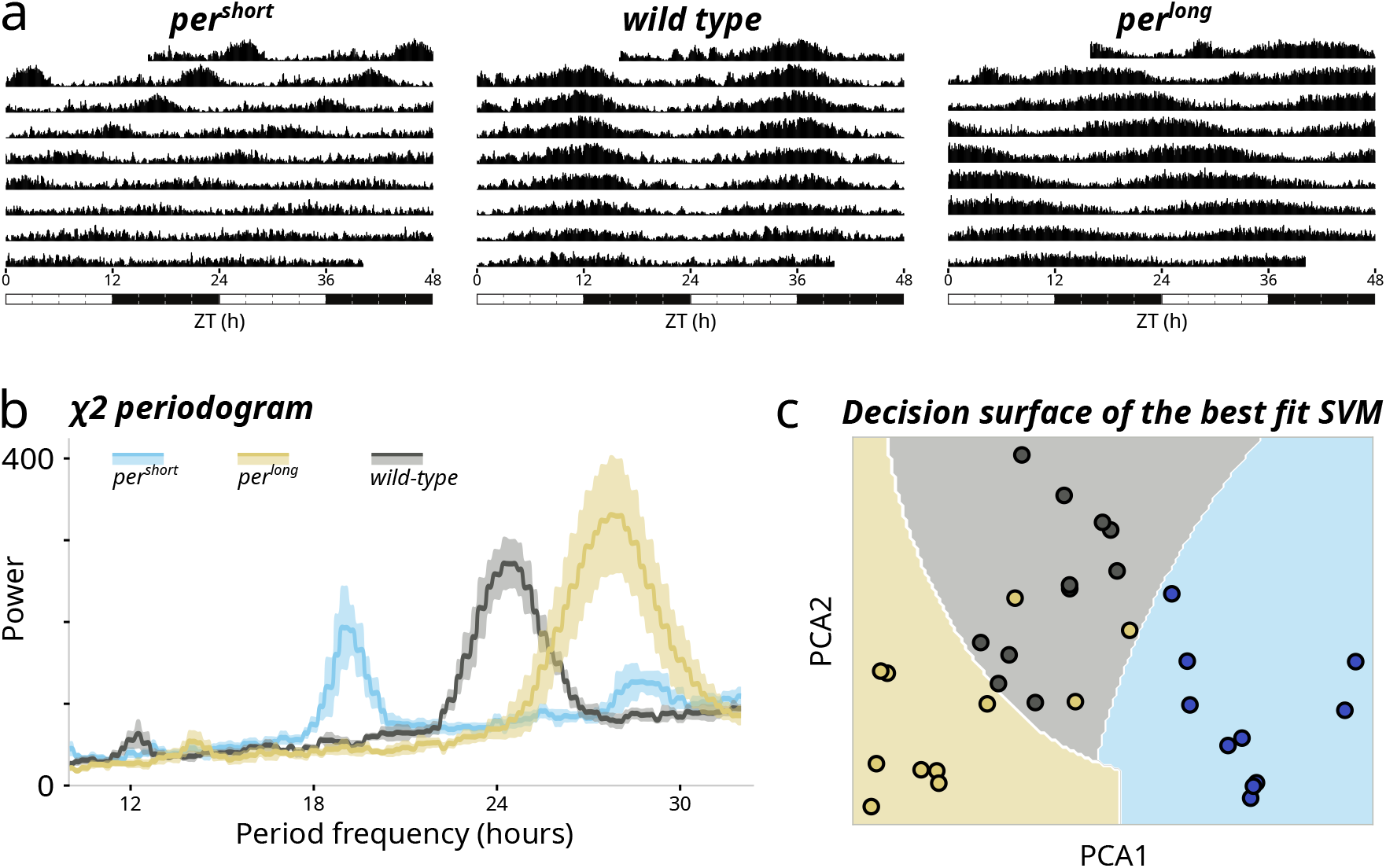
Advanced circadian analysis in ethoscopy. **a)** Double-plotted actograms showing averaged activity during an experiment for animal with short (left), wild-type (middle) or long (right) periods. Time is defined relative to the transition from LD to LL. **b)** Cumulative χ2 periodograms of three populations of animals: short period mutants (*per*^*short*^, light blue), wild type (CantonS, grey), and long period mutants (*per*^*long*^, mustard). **c)** Decision surface (boundaries) of the best fit support vector machine (SVM) separating the data used in b), after performing a catch22 analysis. The three genotype are clustered without supervision. See Notebook 6 in the Supplementary Material for a detailed description of the procedure.

### A first example of ethoscopy new functions: Hidden Markov chain analysis of sleep stages

Wiggin *et al* first proposed an interesting computational paradigm to identify *bona fide* covert sleep stages using Hidden Markov Chains (HMC) (Wiggin *et al*., 2020). Given that their initial analyses were mostly based on data collected using *Drosophila* activity monitors (thus offering only limited resolution of activity (Gilestro, 2012)), we ought to explore how HMC would perform on activity data collected at higher resolution, employing ethoscope video tracking. While we found that the initial sleep characterisation in four hidden sleep stages generally holds true (Figure 1c), we also identified a potentially interesting difference: using ethoscopes, we could detect deep-sleep not throughout the night as originally reported (Wiggin *et al*., 2020), but almost exclusively in the first half of the night, with a clear prominent peak at ZT15 (Figure 1c). Interestingly, this specific window of time was previously shown to be the one associated to the highest arousal threshold using an essay of odour perception (French *et al*., 2021), nicely matching ethoscopy’s description of sleep stages.

### A second example of ethoscopy new functions: Integration with catch22 for unsupervised statistical analysis of time-series

Catch22 (CAnonical Time-series CHaracteristics) is a feature extraction library designed for time-series data analysis (Lubba *et al*., 2019). It provides a set of 22 carefully selected time-series features that capture important characteristics of the data. These features encompass a range of statistical, information-theoretic, and nonlinear measures, making them suitable for a wide variety of time-series applications. Catch22 can be used for feature extraction, comparison of time-series, anomaly detection, classification and regression. Behavioral timeseries are an excellent use case for massive, unbiased features extraction and ethoscope data reveal exceptional discerning power when coupled with these tools (Jones *et al*., 2023). In Notebook 5 and 6 of the Supplementary Material we provide two recipes for analyses on ethoscope. Notebook 5 will guide through integration with HCTSA, a toolbox for massive extraction of more than 7000 features (Fulcher and Jones, 2017), while Notebook 6 will show integration with Catch22, a refinement of the HCTSA set that narrows down to the 22 most commonly informative features (Lubba *et al*., 2019). HCTSA runs in Matlab only, therefore the notebook focuses on bridging ethoscope data to the Matlab code. Catch22, however, features a Python library that well integrates in the ethoscopy pipeline. Notebook 6 and Figure 3c show an implementation of Catch22 combined with classical machine learning algorithms to discern among different circadian phenotypes using unsupervised approaches. This approach can be applied to any phenotype – even when invisible to the human eye – and takes full advantage of Python’s integration with the current machine learning toolbox.

### Conclusions

We built ethoscopy and ethoscope-lab to further improve accessibility and reproducibility in the field of *Drosophila* sleep. Besides offering state-of-the-art data analysis, we envision these tools will open new doors to behavioural scientists and introduce them to good sharing practice in terms of code accessibility and reproducibility.

## Methods

### Software creation

Ethoscopy was written in Python and packaged using Poetry, the Python dependency management and packaging framework. Ethoscopy can be installed using pip (via Pypi) or with the default python distutils. The Dockerfile used to create the ethoscopelab container is shared via github within the ethoscopy repository.

### Data acquisition and analysis

Ethoscope data are collected in single-file SQLite databases (Geissmann *et al*., 2017, 2019) and processed using packages pandas and numpy. By default, sleep is defined as a period of complete inactivity lasting at least five minutes. The hidden Markov chain model is adapted from (Wiggin *et al*., 2020).

## Supporting information

Jupyter Notebooks

## Data availability

The data used to generate the figures and the relative code are available in the tutorial folder of the ethoscopy repository on github (https://github.com/gilestrolab/ethoscopy).

## Code availability

Code, documentation, and tutorials are available at https://lab.gilest.ro/ethoscopy

## Funding and acknowledgements

LB was funded by UKRI with a BBSRC DTP scholarship to Imperial College London (BB/M011178/1 proj 2283755). We thank Esteban Beckwith for providing the circadian data used in one of the Jupyter tutorials and all members of the Gilestro lab for discussion and useful feedback on ethoscopy and ethoscope-lab.

## Author contributions

LB wrote the ethoscopy software with input and guidance from GFG. GFG created the Docker container, wrote the manuscript and supervised the project. Both authors edited the manuscript.

