## Supplementary material for "ethoscopy & ethoscope-lab: a framework for behavioural analysis to lower entrance barrier and aid reproducibility": Jupyter Notebooks: 1_Overview_tutorial.ipynb

```
# %% [markdown]
## Tutorial number 1
### Working with ethoscope data
The tutorial will guide you through loading data, visualisation and plotting using the common sleep functions

# %% [markdown]
## 1. Load the dummy dataset

# %% [code]
import ethoscopy as etho
import pandas as pd

# %% [code]
# This tutorial requires version 1.1.7 or greater
etho.__version__

# %% [code]
# import this function to get the tutorial dataset
from ethoscopy.misc.get_tutorials import get_tutorial

# %% [code]
# There's some dummy data within Ethoscopy for you to play with, use the function below with the argument 'overview' 
# to get the data and metadata that will work with this notebook
# Usually you would have to load the data using the loading functions, head back to the gitbook turotial for information on that
data, metadata = get_tutorial('overview')

# %% [code]
# This creates the behavpy object, combining your data and metadata into a behavpy dataframe (remember to keep them in this order data, metadata - otherwise it will cause problems later down the line)
# The parameter check is used to see if the ids of the specimen in the data match those in the metadata. It also removes some columns that are not needed now you've downloaded the data
# Canvas specifies which plotting library you want to use. The choice is between plotly (default) and the matplotlib-frontend seaborn. 
# plotly allows to output interactive graphs but it's a bit more cumbersome to use and heavier in size. Seaborn / matplotlib is more of a standard, lighter and beautiful but only outputs static plots
# Try this notebook using both canvases to see the differences

df = etho.behavpy(data, metadata, check = True, canvas = 'seaborn')
df.canvas

# %% [code]
# to save on time when looking back on data, save the behavpy dataframe as a pickle file and just load that the subsequent times
df.to_pickle('./tutorial_dataframe.pkl') # replace the path string with your own

# %% [code]
# use this pandas method to load your pickle file
# Once saved and loading below you can skip the steps above
df = pd.read_pickle('./tutorial_dataframe.pkl') # replace with the above

# %% [code]
# this method will display an excerpt of both the metadata and data together for you to view
df.display()

# %% [code]
# you can also just view the data on its own
# python notebooks will print the last line of code without the print function
# If you want to print out of this sequence, use the print function
df
# or print(df)

# %% [code]
# or just the metadata
df.meta

# %% [code]
# this function gives you summary statistics of the whole experiment
df.summary()

# %% [code]
# change the argument of detailed to True for information per sepcimen
df.summary(detailed = True)

# %% [markdown]
## 2. Curate the data

# %% [code]
# Heatmaps a good way to get an overview of the experiment
# you can identify quickly any dead or abnormal specimens
# Ethoscopy uses plotly to generate all data meaning they are interactive and you can zoom by drawing a zooming rectangle 
# Click on the autoscale icon on the top right corner of the image to reset the view to 100% zoom

# Plots will be returned to you from the method/function, its convention to save this variable as fig in each cell
# In the next line juist put fig.show() for the plot to appear below the cell
fig = df.heatmap(variable = 'asleep')
fig.show()

# %% [code]
# calling this method will remove the data for specimens that havent moved for a long time. It retains the data from before the specimen has "died"
df = df.curate_dead_animals()

# %% [code]
# you can filter your data by time to restrict it to periods of interest or to remove data at the start and end which is fragmented
# here we dont need the half data of the first day and anything beyond the day after the sleep deprivation experiment
df = df.t_filter(start_time = 24, end_time = 144)

# %% [code]
# call heatmap again to see the changes done above
fig = df.heatmap(variable = 'asleep')
fig.show()

# %% [code]
# You can also limit your visualisation to a subgroup of animals.
# For instance, we may want to show only animals in the experiment that underwent sleep deprivation
fig = df.xmv('sleep_deprived', True).heatmap( variable = 'asleep')
fig.axes[0].set_title("Sleep Deprived panel in Figure 1a")
fig.show()

# %% [code]
# Or those who did not
fig = df.xmv('sleep_deprived', False).heatmap( variable = 'asleep')
fig.axes[0].set_title("Control panel in Figure 1a")
fig.show()

# %% [code]
# everything looks mostly fine, however there is one specimen - the bottom one on the control group - that appears to be almost always asleep, 
# but has not been removed by the curate_dead_animals() method. This could be just an abnormal specimen or an issue with recording.
# If, after investigation, this turns out to be an artifact or a sick animal, it is OK to remove this specimen.
# Using a notebook like this one will help you keeping track of data curation and data removal. 
# Keeping a log of which animals have to be removed and why is a very important aspect of data handling.

# To do so, zoom in on the row of interest in the heatmap and get the id from the left hand side, for this one it's 2016-04-04_17-39-05_009aee|02
# once we have the id we can remove it with the remove method

df = df.remove('id', '2016-04-04_17-39-05_009aee|02')

# lets call the heatmap once again to see if it's gone
fig = df.heatmap(variable = 'asleep')
fig.show()

# %% [markdown]
## 3. Visualise and save the sleep data

# %% [code]
# The behavpy dataframe has some plotting functions built in to quantify and illustrate your data
# use plot overtime to create an ethogram of your whole dataset
# the solid line is the mean and the shaded area the 95% CI around the mean
fig = df.plot_overtime(variable = 'asleep', title = 'Whole Dataset Ethogram of my first sample experiment')
fig.show()

# %% [code]
# usually you want to compare between experimental groups in your dataset 
# provide the parameter facet_col with a column in metadata to plot each group
# you can also average the data over a day with the parameter wrapped

# Click on the values in the legend to hide / show any particular dataset

fig = df.plot_overtime(variable = 'asleep', wrapped = True, title = 'Ethogram by Experimental group')
fig.show()

# %% [code]
# Sometimes you dont want all the groups from a column in the metdata. To select a subset add a list of the wanted groups to the argument facet_arg
fig = df.plot_overtime(variable = 'asleep', facet_col = 'exp_group', facet_arg = [1,3], title = 'Ethogram by a subset of Experimental group')
fig.show()

# %% [code]
# You can replace the labels in the legend with your own through the facet_labels parameter
# If providing labels you must also provide arguments for facet_arg, as the function cannot guarantee the order will match the labels list
# If your labels or metadata groups contain the words 'baseline', 'control', 'ctrl' then the colour of the plot will be changed to grey automatically
zoomin_df = df.t_filter(start_time=72, end_time=132)
fig = zoomin_df.plot_overtime(variable = 'asleep', facet_col = 'sleep_deprived', facet_arg = [True, False], facet_labels = ['Sleep Deprived', 'Control'], title = 'Ethogram of sleep deprived vs Control - Figure 1b')
fig.show()

# %% [code]
# Sometimes you'll want to split your plots out than more than just one column, 
# 
# If you are using the plotly canvas, you can use the make_tile method
# make_tile() will turn any behavpy plotting method into a subplot
# Here we'll split the dataset by the experimental group and then the sleep deprived group 
# First we have to make the plotting method something we can add as an argument to a method, for which we'll partial()

if df.canvas == 'plotly':
    from functools import partial
    fig_fun = partial(df.plot_overtime, variable = 'asleep', facet_col = 'sleep_deprived', facet_arg = [True, False], facet_labels = ['Sleep Deprived', 'Control'])
    # Then me call the make_tile method with the fig_fun as an argument. The default subplot configuration is 1 colummn and as many rows as in the facet_tile set.
    # However you set your own configuration with the parameters rows and cols, see docstring.

    fig = df.make_tile(facet_tile = 'exp_group', plot_fun = fig_fun)
    fig.show()

    # Currently the make_tile plot can't have the light/dark annotations 
    # It also repeats the same colours per plot 

# This is currently not implemented in seaborn
if df.canvas == 'seaborn':
    pass

# %% [code]
# There are addtional parameters for plot_overtime. 
# 1) Change the moving average window. We usally have the average moving window set to 30 mins, so with 60 second time windows that gives an average window of 30. If you have 10 second bins time window per row
# that means it should be 180. Increasing the number will smooth out the lines of the plot
# 2) Change the bars indicating light / dark phase at the bottom. The default is ZT 12 for lights off, but change it to any integer between 1 and 23 for alternative experiemnts.
# 3) Change the length of a day for the light / dark phase bar with day_length, here we've changed it to 30 hours long with lights off at ZT 15
# 4) Add grid lines from the the y axis to the plot
fig = df.plot_overtime(variable = 'asleep', facet_col = 'sleep_deprived', facet_arg = [True, False], facet_labels = ['Sleep Deprived', 'Control'], title = 'Whole Dataset Ethogram with changes', avg_window = 180, day_length = 30, lights_off = 15, grids = True)
fig.show()

# %% [code]
# You can save your plots as image files, pdfs, and as html files
# saving as a html file retains plotys interactive attribrutes and is best for when you want to view your plot on the whole screen and not just a small notebook output
# to save call the function below on your returned fig object
# Remember to change the location to the folder you are working in
fig = df.plot_overtime(variable = 'asleep', facet_col = 'exp_group')

etho.save_figure(fig, './tutorial_plot.html')
# for non html files you can specify the width and height of the saved figure
etho.save_figure(fig, './tutorial_plot.pdf', width = 1500, height = 1500)
etho.save_figure(fig, './tutorial_plot.png', width = 2000, height = 1500)

# %% [code]
# Quantify your ethograms with plot_quantify
# This method takes the same parameters as plot_overtime (except avg_window and circadian_night and wrapped)
# Here we want to look at the baseline sleep so we filter it to before the experimental day
# hover on the plot with your mouse to get the mean and 95% confidence intervals around it, as well as individual specimens values
# Every quantifying plot returns a data frame with the plotted values per group for further statistical analysis

temp_df = df.t_filter(start_time = 24, end_time = 96)
fig, stats_quant = temp_df.plot_quantify(variable = 'asleep', facet_col = 'sleep_deprived', facet_arg = [True, False], facet_labels = ['Sleep Deprived', 'Control'], title = 'Quantification of Basline Sleep')
fig.show()

# %% [code]
# The returned pandas dataframe has the mean values per experimental group
# Row colunms are not related to each other, each column is a seperate list
stats_quant

# %% [markdown]
## 4. Quantify and compare

# %% [code]
# Here we want to look at the the rebound period (3 hours post) after sleep deprivation
temp_df = df.t_filter(start_time = 120, end_time = 123)
fig, stats_deprivation = temp_df.plot_quantify(variable = 'asleep', facet_col = 'sleep_deprived', facet_arg = [True, False], facet_labels = ['Sleep Deprived', 'Control'], title = 'Quantification of Rebound Sleep\nFigure 1c')
fig.show()

# %% [code]
# Often you'll want to seperate day and night data to see differences between the phases
# Like plot_quantify, plot_day_night also takes the same parameters
temp_df = df.t_filter(start_time = 24, end_time = 96)
fig, stats_day_night = temp_df.plot_day_night(variable = 'asleep', facet_col = 'sleep_deprived', facet_arg = [True, False], facet_labels = ['Sleep Deprived', 'Control'], title = 'Quantification of Day and Night')
fig.show()

# %% [code]
# Sometimes you'll want to compare two or more variables from the data
# The ethoscope can detect movement at a micro movmement level, differences in the ratios of micromovment to walking can help further describe a specimen
# Compare varaibles has two y-axes, the first variable of the list will use the left y-axis and the last in the list will use the right y-axis. Any in the middle will use the left, so list accordingly
temp_df = df.t_filter(start_time = 24, end_time = 96)
#fig, stats_multiple = temp_df.plot_compare_variables(variables = ['micro', 'walk', 'max_velocity'], facet_col = 'sleep_deprived', facet_arg = [True, False], facet_labels = ['Sleep Deprived', 'Control'], title = 'Quantification of Micro movements Vs Walking')
fig, stats_multiple = temp_df.plot_quantify(variable = ['micro', 'walk', 'max_velocity'], facet_col = 'sleep_deprived', facet_arg = [True, False], facet_labels = ['Sleep Deprived', 'Control'], title = 'Quantification of Micro movements Vs Walking')
fig.show()

# %% [markdown]
## 5. Altering figures before show()

# %% [code]
# As you have the fig object you can edit all its attributes, so if you don' like our axis titles you can change them
# The way you do this, depends on the type of canvas you want to use


fig = df.plot_overtime(variable = 'asleep', facet_col = 'exp_group')

if df.canvas == 'plotly':
# Here's a few basic examples for the plotly canvas but head to https://plotly.com/python/reference/ for examples on what can be changed 

    # let's change the x-axis
    fig['layout']['xaxis']['title'].update(text = 'new x-axis label', font = dict(color = 'red', size = 30))
    # Now the y-axis
    fig['layout']['yaxis']['title'].update(text = 'new y-axis label', font = dict(color = 'blue', size = 10))
    fig.show()

if df.canvas == 'seaborn':
# And this is the same done in seaborn, which is based on matplotlib https://seaborn.pydata.org/

    fig.axes[0].set_xlabel('new x-axis label', color='red', fontsize=30)
    fig.axes[0].set_ylabel('new y-axis label', color='blue', fontsize=30)


# %% [code]
fig = df.plot_overtime(variable = 'asleep', facet_col = 'exp_group')

# let's change the range and the position of the ticks

if df.canvas == 'plotly':

    fig['layout']['yaxis'].update(range = [0,2], dtick = 0.5)
    fig.show()

elif df.canvas == 'seaborn':

    import numpy as np

    fig.axes[0].set_ylim([0, 2])    # Set the limits of y-axis
    fig.axes[0].yaxis.set_ticks(np.arange(0, 2.1, 0.5))    # Set tick locations


# %% [markdown]
## 6. Export the notebook for your publications

# %% [code]
# Notebooks are great for record keeping and good practice sharing
# Export this entire notebook as HTML so that anyone can have access to your code and the figures it generates without even having to install python

# File -> Save and Export Notebook As -> HTML
```
