## Supplementary material for "ethoscopy & ethoscope-lab: a framework for behavioural analysis to lower entrance barrier and aid reproducibility": Jupyter Notebooks: 2_HMM_tutorial.ipynb

```
# %% [markdown]
## Tutorial number 2
### Identify sleep stages with Hidden Markov Models

# %% [markdown]
This tutorial assumes you have some basic knowledge of Ethoscopy, if you don't please head to the overview tutorial first.

For this tutorial we will be training a Hidden Markov Model (HMM) using an ethoscopy wrapper class for the python package Hmmlearn, see https://hmmlearn.readthedocs.io/en/latest/ for more details on what this excellent package can do. A good introduction to HMM's is available here: https://web.stanford.edu/~jurafsky/slp3/A.pdf

Ethoscopy adapts their Categorical HMM, as such the model can only be trained using discrete data represented as integers starting with 0 as the first. I.e. "Cat 1 = 0, Cat 2 = 1, ..."

At the Gilestro lab we train our model using movement data from the Ethoscope system, using tested parameter limits for velocity as movement we categorise movement as either True or False, or 1 or 0. During the setup we found data binned to 60 seconds (1 min) and then trained best recapitulated the sleep of CS drosophila.

# %% [markdown]
## 1. Load the dummy dataset

# %% [code]
import pickle
import ethoscopy as etho
import numpy as np

# %% [code]
# This tutorial required version 1.1.8 or greater
etho.__version__

# %% [code]
# import this function to get the tutorial dataset
from ethoscopy.misc.get_tutorials import get_tutorial

# %% [code]
# We'll be using the same dataset as in the overview tutorial, so use the same function below with the argument 'overview' 
# Load the data and metadata, and then intialise it into a behavpy_HMM 
data, metadata = get_tutorial('overview')
df = etho.behavpy_HMM(data, metadata, check = True)

# %% [markdown]
## 2. Curate the Data

# %% [code]
# First we'll call the method to remove dead specimens
df = df.curate_dead_animals()
# We'll also remove the specimen we noticed in the overview tutorial that was missed by the above method
df = df.remove('id', '2016-04-04_17-39-05_009aee|02')

# %% [code]
# As we saw in the overview tutorial the dataset we have has half the specimens sleep deprived later in the experiment
fig = df.heatmap(variable = 'asleep')
fig.show()

# %% [code]
# When training a HMM you want a specimens basline data as normal conditions, so for this excerise we'll want to trim the dataset
# We can see from the heatmap that the sleep deprivation begins at 108 hours (ZT 12), we'll also trim the first day data as the specimens settle in
df = df.t_filter(start_time = 24, end_time = 108)

# %% [code]
# It's also key to have as few missing values as possible when training the data, as the model assumes a consitent time gap between each point
# Our dataset now has 3.5 days with time bins of 60 seconds, so if a specimen was to have maximum data points for that time they would have
# (86400/60) * 3.5 rows of data. Lets not take any chances and only take specimens with that exact amount
df = df.curate(points = (86400/60) * 3.5)

# %% [code]
# If any of the data passed to the model to be trained contains NaN values it will throw an error
# So lets check our data for NaN values
df['moving'][df['moving'].isnull()]

# %% [code]
# We have one NaN value, which is enough to crash the whole thing!
# Luckily pandas has built in methods to replace NaN values, using the method below with "ffill" fills in the NaN with the value of the row above (previous)
df['moving'] = df['moving'].fillna(method = "ffill")

# Lets to see if it worked
df['moving'][df['moving'].isnull()]

# %% [markdown]
## 3. Setup

# %% [code]
# Name the hidden states you want to fit the model to - at the Gilestro lab we use a 4 state model, but feel free to play around!
hidden_states = ['Deep sleep', 'Light sleep', 'Light awake', 'Active awake']

# Declare your observables, the first will equal the value 0 in your data
observables = ['Inactive', 'Active']

# This is key to training your model, where you declare the interaction between the hidden states
# The array shape must be the length of your states_list by itself, E.g. 4x4 for the above
# The first nested array equals the first in your list, with each index relative to it's list position.
# E.g. For Deep Sleep (the first array), we have that it can transition into all other states and itself other than Full Awake (the last position)
# For Active awake (the third array), it can't transtion into Deep Sleep (the first position), but can transition into everything else and itself
# If you want a state to tranition into another we recommend you have it as 'rand', this randomises the starting transtion rate before the model iterates and updates.
# This is so we can run the training multiple times to then compare scores, hopefully minimising the risk of getting into a local minima
# If you wish to declare the starting transtions do so as a float with each nested array totalling 1.0 and set iterations to 1
t_prob = np.array([['rand', 'rand', 'rand', 0.0],
                    ['rand', 'rand', 'rand', 0.0],
                    [0.0, 'rand', 'rand', 'rand'],
                    [0.0, 0.0, 'rand', 'rand']])

# Here we do the same, but for the emmission probabilites (liklihood a state will be observed as one of the observable states)
# The same rules for array shape apply, but for the observable list
# Here the first two states are sleep states, so we declare that they can't be active during that time
em_prob =  np.array([[1.0, 0.0],
                    [1.0, 0.0],
                    ['rand', 'rand'],
                    ['rand', 'rand']])

# We also need to changed the boolean values of 'moving' into integers for the model to work
# np.where will replace all values where the argument is True with the first argument, all those that are false with the second
df['moving'] = np.where(df['moving'] == True, 1, 0)

# %% [code]
# Now we're setup we can call the training method
# iterations is the number of times you want to iterate and train starting with new transiton and emission probabilities. It's recommended to have this higher than 3
# but for the tutorial we'll keep it low for the sake of time. More iterations mean you're more likely to find the best fitting model and
# hmm_iterations is the number of times the HMM will iterate with newer parameters before stopping
# tol is the the value for the difference in log liklihood after each iteration, once the value equals your input the model will stop updating the parameters
# After each time the transiton and emission probabilities are randomised and a model is produced, a score of logliklihood will be generated against 10% of the data,
# each new model will be scored against this and the best one will be returned at the end. Change the size of the test batch with test_size
# Save the best model to your local as a pickle file with file_name. The method will also return the trained model, but it's best to save it to a file

h = df.hmm_train(
    states = hidden_states, 
    observables = observables, 
    var_column = 'moving', 
    trans_probs = t_prob, 
    emiss_probs = em_prob, 
    start_probs = None, 
    iterations = 3, 
    hmm_iterations = 100, 
    tol = 2, 
    t_column = 't', 
    bin_time = 60, 
    test_size = 10,
    file_name = 'tutorial_hmm.pkl', # replace with your own file name. If you already have a pickle file named this, it will test the first iteration against it.
    verbose = False)

# If verbose is True it will print to screen the logliklihood score of each hmm iteration

# %% [code]
# if .hmm_train() has an argument for the file_name parameter a pickle copy of the trained HMM will be saved to your local
# Use the below to load the trained model for analysis
with open('./tutorial_hmm.pkl', 'rb') as file: 
    h = pickle.load(file)

# %% [markdown]
## 4. Plotting the decoded data

# %% [code]
# Lets have a look at our trained HMM
# Here we can see the transition rates between the difference states
df.hmm_display(hmm = h, states = hidden_states, observables = observables)

# %% [code]
# Ethoscopy also has pretrained HMMs for both Male and Female CS Drosophila, trained on several days worth of data on hundreds of flies
# These have the state structure of the above
# Feel free to use these when looking at your own wildtype datasets, however if using other species or mutants you will need to train your own
# For the rest of this tutorial we'll use a the Male pretrained model

# first we need to import the get_HMM function
from ethoscopy.misc.get_HMM import get_HMM

# %% [code]
# Have the argument as "M" or "F" for the male or female dataset respectively
h = get_HMM("M")

# %% [code]
# Plotting the specimens HMM overtime is much like plot_overtime(), except you cannot facet by the metadata, see split plots below!
# The only addtional arguments are for the trained HMM, the label names for each state and their colours, and also the time each row is binned when it was trained, i.e. 60 seconds (default)
# If you wish to follow our naming and colour schema then leave labels and colours as None, if you have a 4 state model it will use the labels as seen above and our blue/red colour scheme
# You can change day_length, lights_off, title, grids as seen previously
# Save plots giving an arugment to parameter save with the location and file type to be saved as, see overview tutorial for more information
fig = df.plot_hmm_overtime(
            hmm = h, 
            variable = 'moving', 
            wrapped = True, 
            bin = 60, 
            title = 'A decoded HMM over one day - Figure 2d'
            )
fig.show()
etho.save_figure(fig, "./figure2d.svg")

# %% [code]
# Here's it with new names and different colours
# The colours must follow Plotlys colour naming convention
fig = df.plot_hmm_overtime(
            hmm = h, 
            variable = 'moving', 
            labels = ['Out Cold', 'Soft Sleep', 'Dozing', 'On One'], 
            colours = ['black', 'yellow', 'pink', 'red'],
            wrapped = True, 
            title = 'Your own Colour Scheme'
)
fig.show()

# %% [code]
df.groupby('id').count()

# %% [code]
# Due to having multiple states plotted at once its easier to compare by faceting with multiple plots for each states
# plot_hmm_split is the same as plot_hmm_overtime, but with the facet_labels, facet_col, facet_arg parameters we've seen before to compare via our metadata
fig = df.plot_hmm_split(
            hmm = h, 
            variable = 'moving', 
            facet_col = 'exp_group', 
            wrapped = True, 
            bin = 60, 
            title = "Split plot per Experimental Group"
)
fig.show()

# %% [code]
# To quantify the above just call the plot_hmm_quantify method 
# Like all the quantify methods it will also return a dataframe with the values of each plotted dot for you to do any statistical analysis
# The plotted and returned values have all values with a z-score > 3/-3 removed
fig, stats_time = df.plot_hmm_quantify(
            hmm = h, 
            variable = 'moving', 
            facet_col = 'exp_group', 
            title = "Quantify each State"
)
fig.show()

# %% [code]
# To quantify the above just call the plot_hmm_quantify method 
# Like all the quantify methods it will also return a dataframe with the values of each plotted dot for you to do any statistical analysis
# The plotted and returned values have all values with a z-score > 3/-3 removed
fig, stats_time = df.plot_hmm_quantify(
            hmm = h, 
            variable = 'moving', 
            title = "Quantify each State - Figure 2e"
)
fig.show()
etho.save_figure(fig, "./figure2e.svg")

# %% [code]
# To quantify the above just call the plot_hmm_quantify method 
# Like all the quantify methods it will also return a dataframe with the values of each plotted dot for you to do any statistical analysis
# The plotted and returned values have all values with a z-score > 3/-3 removed
fig, stats_time = df.plot_hmm_quantify(
            hmm = h, 
            variable = 'moving', 
            title = "Quantify each State"
)
fig.show()

# %% [code]
# The average is useful, but it doesn't tell us about about how time influences the decision by the model to predict each state
# Use plot_hmm_quantify_length_min_max to see the maximum and mimimum length of each state to understand when the model switches it's predictions
# Here we can see that light sleep can only be 11 minutes of inactivity long, after that it is predicted as deep sleep
fig = df.plot_hmm_quantify_length_min_max(
    hmm = h, 
    variable = 'moving', 
    title = "The Max and Min length of each state"
)
fig.show()


# %% [code]
# It is also useful to see how often a state is transitioned into, to understand its frequency in relation to length
# Here we can see the two light sleep and awake stages are transitioned into often, which compliments their low run lengths
# In contrast the other two states are transitioned into less frequently, but when they do the it is for a lot longer
fig, stats_transition = df.plot_hmm_quantify_transition(
    hmm = h, 
    variable = 'moving', 
    title = "Quantify the average transtion of each State"
)
fig.show()

# %% [code]
# All of the above (except plot_hmm_overtime) can take arguments of multiple trained HMMs with the view that can compare different species, mutants, ect.
# For example to demonstrate we can take this dataset and compare two groups, one using the male trained model and the other the female trained model
# When you have multiple HMMs you must have the same number of arguments, labels, and bins. Otherwise you will raise an error
# Try this out with the other plotting functions!
h_f = get_HMM("F")
fig = df.plot_hmm_split(
    hmm = [h, h_f], 
    variable = 'moving', 
    facet_col = 'exp_group', 
    facet_arg = [1, 2], 
    facet_labels = ['Male', 'Female'], 
    wrapped = True, 
    bin = [60, 60], 
    title = "Split plot with two different HMMs"
    )
fig.show()

# %% [code]
# Finally, sometimes you want to see the decoded data minute by minute to get to understand it properly
# Like the others you can have multiple HMMs, if you choose to do this then it will display the same specimen but decoded by each HMM
# The ID of each specimen will be printed to screen in case you want to investigate further as each ID is picked randomly each time it's run
fig = df.plot_hmm_raw(
    hmm = h, 
    variable = 'moving', 
    num_plots = 5, # the number of different specimens you want in the plot 
    bin = 60, 
    title = ''
)
fig.show()

# Addtionally, if you have response data from mAGO experiments you can add that behavpy dataframe to highlight points of interaction and their response 
# Purple = response, lime = no response
## df.plot_hmm_raw(mago_df = mdf, hmm = h, variable = 'moving', num_plots = 5, bin = 60, title = '', save = False)

# %% [code]
```
