## Supplementary material for "ethoscopy & ethoscope-lab: a framework for behavioural analysis to lower entrance barrier and aid reproducibility": Jupyter Notebooks: 3_Circadian_tutorial.ipynb

```
# %% [markdown]
## Tutorial number 3
### Circadian analysis on ethoscope data

# %% [markdown]
This tutorial assumes you have some basic knowledge of Ethoscopy, if you don't please head to the overview tutorial first.

This tutorial will walk through the array of circadian analysis methods ethoscopy contains, from actograms and anticipation scores, to periodograms. Whilst the tutorial will keep things basic there will be some assumed knowlege about circadian rythmn.

# %% [markdown]
## 1. Load the dummy dataset

# %% [code]
import ethoscopy as etho

# %% [code]
# This tutorial required version 1.2.0 or greater
etho.__version__

# %% [code]
# import this function to get the tutorial dataset
from ethoscopy.misc.get_tutorials import get_tutorial

# %% [code]
# We'll be using a circadian specific dataset this time that was generated previously, enter 'circadian' as the argument to grab it
# Load the data and metadata, and then intialise it into a behavpy object
data, metadata = get_tutorial('circadian')
df = etho.behavpy(data, metadata, check = True, canvas="plotly")
df.canvas

# %% [markdown]
## 2. Looking at the data

# %% [code]
# Lets have a look at what we're working with
df.summary()

# %% [code]
# In the meta we can see the specimens are split into different experimental groups, these are the groups we will want to look at
df.meta

# %% [code]
# We can see from plotting the movement data that they a different rythmn
fig = df.plot_overtime('moving', facet_col = 'period_group')
fig.show()

# %% [markdown]
## 3. Actograms

# %% [markdown]
Actograms are a simple visual way to see changes in activity over multiple days. With ethoscopy you have two ways of viewing your actograms, you can create one for every specimen or you can average it over the whole dataframe / experimental group

# %% [markdown]
### Grouped Actograms

# %% [code]
# Lets create an averaged Actogram for each period group using the .plot_actogram() method
# Choose the column with the movement data, the averaging window you'd like to use, and then the column to facet by
fig = df.plot_actogram(
        mov_variable = 'moving',
        bin_window = 5, # the default is 5, but feel free to change it to smooth out the plot or vice versa
        facet_col = 'period_group',
        facet_arg = ['short', 'wt', 'long'],
        title = 'Grouped Actogram',
        #figsize = (16,4)
        )

fig.show()

# If you dont give an arugment to facet_col then a single actogram will be plotted that is an average of all your data

# %% [markdown]
### Tiled Actograms

# %% [code]
# Although we can already see the differences between the groups, often when you don't already have a nicely curated dataset it's nice to see everything
# When plotting all specimens the default is to take the specimens IDs as each plots title. However, these names can be uniformative at first glance and we might want
# to replace them with more informative labels. Lets create some for this example

# Remember to access the metadata part of the dataframe just add .meta
# We can create a new column by combining two others. However, the regio_id is a integer and not a string, so we have to convert it first
df.meta['tile_labels'] = df.meta['period_group'] + '_' + df.meta['region_id'].apply(lambda l: str(l))

# %% [code]
# plot_actogram_tile will plot every specimen in your behavpy dataframe

if df.canvas == "plotly":
        fig = df.plot_actogram_tile(
                mov_variable = 'moving',
                labels = 'tile_labels',
                title = 'Tiled Actogram')

        fig.show()

        # This is a good example of a plot that is hard to view inside a jupyter notebook, so lets save it as a html file and view it in our browser in full screen
        etho.save_figure(fig, './tiled_actogram.html')
        # right click the saved file and click open in browser

elif df.canvas == "seaborn":
        fig = df.plot_actogram(
        mov_variable = 'moving',
        bin_window = 5, # the default is 5, but feel free to change it to smooth out the plot or vice versa
        facet_col = 'tile_labels',
        title = 'Tiled Actogram',
        figsize=(60,30))

        fig.show()

# %% [code]
# If you just want to see produce a single actogram for one specimen then filter the dataframe before and then call .plot_actogram()
temp_df = df.xmv('id', '2017-01-16 08:00:00|circadian.txt|01')

fig = temp_df.plot_actogram(
        mov_variable = 'moving',
        title = 'Individual Actogram - Figure 3a',
        #figsize=(12,4)
        )

fig.show()
etho.save_figure(fig, "./figure3a.svg")

# %% [markdown]
## 4. Anticipation Score

# %% [markdown]
The anticipation score is the ratio of the final 3 hours of activity compared to the whole 6 hours of activity prior to lights on/off. This gives us a way of quantifying the morning and evening peaks of activity that are hallmarks of Drosophila circadian rythmn.

# %% [code]
# Simply call the plot_anticipation_score() method to get a box plot of your results
# the day length and lights on/off time is 24 and 12 respectively, but these can be changed if you're augmenting the environment

fig, stats = df.plot_anticipation_score(
        mov_variable = 'moving', 
        facet_col = 'period_group',
        facet_arg = ['short', 'wt', 'long'],
        day_length = 24,
        lights_off = 12,
        title = 'Anticipation Score')

fig.show()

# Here we can see only the WT flies have over 50% of their activity in the final 3 hours before lights off. This shows that they are anticipating the evening feeding time correctly,
# whereas the others fail to do so. Lights On score is the same across the groups.

# %% [markdown]
## 4. Periodograms

# %% [markdown]
Periodograms are essential for definitely showing periodicity in a quantifiable way. Periodograms often make use of algorithms created from spectral analysis, to decompose a signal into its component waves of varying frequencies. This has been adopted to behavioural data, in that it can find a base rythmn over several days from what is usually unclean data.

Ethoscopy has 5 types of periodograms built into its behavpy_periodogram class, which are 'chi squared' (the most commonly used), 'lomb scargle', fourier, and 'welch' (all based of of the fast fourier transformation (FFT) algorithm) and 'wavelet' (using FFT but maintaining the time dimension). 

Try them all out on your data and see which works best for you.

# %% [code]
# To access the periodogram methods we first have to change the behavpy object
per_df = etho.behavpy_periodogram(df, df.meta, check = True)

# %% [code]
# Before we can access all the methods we first must compute our periodogram, so always call this method first
# For this example we'll compare the chi squared periodogram with lomb scargle
# the periodogram can only compute 4 types of periodograms currently, choose from this list ['chi_squared', 'lomb_scargle', 'fourier', 'welch']
per_chi = per_df.periodogram(
        mov_variable = 'moving',
        periodogram = 'chi_squared', 
        period_range = [10,32], # the range in time (hours) we think the circadian frequency lies, [10,32] is the default
        sampling_rate = 15, # the time in minutes you wish to smooth the data to. This method will interpolate missing results
        alpha = 0.01 # the cutoff value for significance, i.e. 1% confidence
)

per_lomb = per_df.periodogram(
        mov_variable = 'moving',
        periodogram = 'lomb_scargle', 
        period_range = [10,32],
        sampling_rate = 15,
        alpha = 0.01 
)

# Hint! Sometimes increasing the sampling_rate can improve you're periodograms. On average we found 15 to work best with our data, but it can be different for everyone.

# %% [code]
# each periodogram should have atleast the columns period, power, and sig_threshold
per_chi

# %% [code]
# We can see that lomb_scargle has more rows that chi_squared. This is because the loomb scargle returns a list of freqeuncies between our period range that it determines best represents the
# data given. However, chi squared will just binned windows we give it from the sampling_rate
# Lomb scargle has been reported to be more accurate with chi squared sometimes giving false positives, see: https://journals.plos.org/ploscompbiol/article?id=10.1371/journal.pcbi.1008567
# However sometimes it's the only one that works, so try all of them
per_lomb

# %% [markdown]
### Plotting the Periodograms

# %% [markdown]
#### Tiled Plots

# %% [code]
# To get a good understanding of each individual specimen you might want to plot the periodograms of each individual
# Like the actogram_tile method we can rely on the ids of each specimen or we can use the labels we created
fig = per_chi.plot_periodogram_tile(
            labels = 'tile_labels',
            find_peaks = True, # make find_peaks True to add a marker over significant peaks
            title = 'Tiled Chi Squared Periodogram'
)
fig.show()

# %% [code]
fig = per_lomb.plot_periodogram_tile(
            labels = 'tile_labels',
            find_peaks = True, # make find_peaks True to add a marker over significant peaks
            title = 'Tiled Lomb Scargle Periodogram'
)
fig.show()

# %% [markdown]
#### Grouped Plots

# %% [code]
# Lets have a looked at them averaged across the groups
fig = per_chi.plot_periodogram(
            facet_col = 'period_group',
            facet_arg = ['short', 'wt', 'long'],
            title = 'Grouped Chi Squared Periodogram - Figure 3b'
)
fig.show()

etho.save_figure(fig, "./figure3b.svg")

# %% [code]
fig = per_lomb.plot_periodogram(
            facet_col = 'period_group',
            facet_arg = ['short', 'wt', 'long'],
            title = 'Grouped Lomb Scargle Periodogram'
)
fig.show()

# %% [markdown]
We can see from both Chi Squared and Lomb Scargle that the different groups have periodicities of about 19, 24, and 27 respectively. But now we want to quantify this.

# %% [markdown]
#### Quantify Periodogram

# %% [code]
# Quantify the periodograms by finding the highest power peak and comparing it across specimens in a group
# call .quantify_periodogram() to get the results
fig, stats = per_chi.quantify_periodogram(
        facet_col = 'period_group',
        facet_arg = ['short', 'wt', 'long'],
        title = 'Quantification of Chi Squared Periodogram'
)
fig.show()

# %% [code]
# Ethoscopy also has pretrained HMMs for both Male and Female CS Drosophila, trained on several days worth of data on hundreds of flies
fig, stats = per_chi.quantify_periodogram(
        facet_col = 'period_group',
        facet_arg = ['short', 'wt', 'long'],
        title = 'Quantification of Chi Squared Periodogram'
)
fig.show()

# %% [code]
# You can find the peaks in your periodograms without plotting, use .find_peaks()
# Find peaks adds a column to your periodogram dataframe with the rank of your peak. I.e. false = no peak, 2 = 2nd highest peak, 1 = highest peak
per_chi_peaks = per_chi.find_peaks(
            num_peaks = 3 # maximum number of peaks you want returned, if there are less than your given number then only they will be returned
)
#Lets have a look at the peaks
per_chi_peaks[per_chi_peaks['peak'] != False]

# %% [markdown]
### Wavelets

# %% [markdown]
The final periodogram is wavelets, which preserve the time dimension when finding the underlying frequencies. This means you can see how periodcity changes over the course of a long experiment

# %% [code]
# Wavelet plots are intensive to plot usually due to the amount of points. Therefore the .wavelet function will compute and plot just one wavelet transformation, averaging all the data in your dataframe
# It's therefore recommended you filter prior to calling the function, either by groups or to individual specimens
# This method is powered by the python package pywt, Head to https://pywavelets.readthedocs.io/en/latest/ for information about the pacakage and the other wavelet types
wt_df = per_df.xmv('period_group', 'wt')
fig = wt_df.wavelet(
        mov_variable = 'moving',
        sampling_rate = 15, # increasing the sampling rate will increase the resolution of lower frequencues (16-30 hours), but lose resolution of higher frequencies (2-8 hours)
        scale = 156, # the defualt is 156, increasing this number will increase the resolution of lower frequencies (and lower the high frequencies), but it will take longer to compute
        wavelet_type = 'morl', # the default wavelet type and the most commonly used in circadian analysis. Head to the website above for the other wavelet types or call the method .wavelet_types() to see their names
        title = 'Wavelet: WT'

)
fig.show()

# %% [markdown]
Areas of organge/red have high scores for the freqencies on the y-axis. Due to trying to fit all of the frequencies of the y-axis in the results are plotted on a natural log scale, this makes it hard to find accurate changes in periodicity. However we can see how it changes over time, which here seems to decrease slightly. (The edge cases of wavelets tend to have the lowest scores due to the effect of windowing, so the drop is not unexpected)

# %% [markdown]
Here we've walked you through how to use behavpy to perform circadian analysis on your data, go forth and analyse
```
