## Supplementary material for "ethoscopy & ethoscope-lab: a framework for behavioural analysis to lower entrance barrier and aid reproducibility": Jupyter Notebooks: 4_Navigating_db_tutorial.ipynb

````
# %% [markdown]
## Tutorial number 4
### Navigating the ethoscope db
ethoscopy provides some accessory functions to navigate your local ethoscope repository and sort SQLlite3 db files.
This is particularly handy post-publication, when db files linked to an experiment need to be identified and shared 
with the community

# %% [code]
from ethoscopy.metadata_db import db_organiser, metadata_handler, db_crawler

# %% [code]

# This is the step in which we crawl through our folder containing all the db files from the lab
# If refresh is set to True, it will scan all the files and generate a new csv list of all the files available in the system
# If refresh is set to False, it will simply load a pre-compiled csv list (like in the case of this tutorial)
# We provide a copy of `ethoscope_db.csv` which list all the 16.8k db files of the Gilestro lab as of Dec 2022

datapath = "/mnt/ethoscope_results"
db = db_organiser(datapath, refresh=False)

# %% [code]
# Now we load the metadata file and do the first processing 
# to try and understand how many db files we should be looking for
# This is a real dataset from Jones_et_al 2023

metadata_filename = 'jones_et_al_metadata.csv'
meta = metadata_handler(metadata_filename, project='coccinella_2022', authors=['Jones'] )
meta.db_files

# %% [code]
# This is where we go through the metadata and find the relevant db files
meta.associate_to_db(db)
meta.summary

# %% [code]
# It seems we have 8720 in our metadata list, corresponding to 381 files
# However, we could not find 3 of these 381, corresponding to 72 animals
# To see which experiments are missing (probably because they were mislabeled) use the following:
meta.list_dbs(notfound = True)

# %% [code]
# Which can be saved in the following way
meta.list_dbs(notfound = True).to_csv('jones_et_al_dbs_not_found.txt')

# %% [code]
# Conversely, if we want to write all the other succesfully found db to a csv we can use the following:
meta.list_dbs().to_csv('jones_et_al_all_info.txt')

# %% [code]
# Those options however are rather verbose, providing also info about the db filesize and so on
# IF we want only the filenames and location of each db we can use
meta.list_dbs().db_filename.to_csv('jones_et_al_only_dbs.txt', header=None, index=None)

# %% [markdown]
### Upload all the db files to Zenodo

# %% [markdown]
Now we have a handy list of all the db files associated to our experiments, ready to be shared with the world.
Notice that these 381 files take 168.73 GB which is above the 50GB limit that we would have on a Zenodo repository.
We will have to zip them and use three repositories. **The following is not Python but bash!**

# %% [markdown]
First we need to copy all these files to a temporary destination
The following will create a copy of all the files we want to upload in the repo

```
mkdir /mnt/data/temporary/jones_et_al
rsync -aruRP --files-from=jones_et_al_only_dbs.txt /mnt/data/results /mnt/data/temporary/jones_et_al/
```

# %% [markdown]
Now we zip all these files in multiple archives, each 2GB in size
```
zip -r -s 2000m jones_et_al_db_files.zip jones_et_al
```

# %% [markdown]
Now create some descriptive files of what is in the archive
```
cd /mnt/data/temporary/jones_et_al
```
create a txt file providing the md5sum of each zip archive
```
find . -type f -name "*.z*" -exec md5sum {} + > ../all_zips_with_checksum.txt
```

same thing but for the original uncompressed db files
```
find . -type f -name "*.db" -exec md5sum {} + > ../all_dbs_with_checksum.txt
```

visualize the structure of the folders / files
```
tree > jones_et_al/db_files_content.txt
```

# %% [code]
````
