## Supplementary material for "ethoscopy & ethoscope-lab: a framework for behavioural analysis to lower entrance barrier and aid reproducibility": Jupyter Notebooks: 5_Ethoscopy_to_hctsa_tutorial.ipynb

```
# %% [markdown]
## Tutorial number 5
### Exporting ethoscope data to `.mat` format for integration with HCTSA

# %% [markdown]
This tutorial will take you though converting a behavpy dataset into a format that can be input into HCTSA, a time series comparative analysis software. Currently the full suite of software only works in Matlab. So this tutorial shows you have to convert the data into .mat file that you can then use in the HCTSA environment. The .mat file will include the time series data per specimen, the labels (id), and keywords per specimen.

If you want to just stick with Python see tutorial 5 which uses HCTSA smaller toolbox catch22, limited to the top 22 most common analytical methods from HCTSA, fully in a Python environment. The use of Catch22 is reccomended.

# %% [markdown]
## 1. Load the dummy dataset

# %% [code]
import ethoscopy as etho
import pandas as pd
import numpy as np
from functools import partial
# we need the scipy function savemat to create a .mat file
from scipy.io import savemat

# %% [code]
# Load in the data from the overview tutorial
from ethoscopy.misc.get_tutorials import get_tutorial
data, metadata = get_tutorial('overview')
df = etho.behavpy(data, metadata, check = True)

# %% [markdown]
## 2. Choose the variable you want to analyse in HCTSA

# %% [code]
# Lets use x (the position of the fly) for this tutorial
var = 'x'

# %% [markdown]
### HCTSA needs time series data of the same length so we need to do some data curation

# %% [code]
# The ethoscope if loading with sleep_annoation will interpolate data about sleep, this causes NaN values for the normal variables other than 'asleep'
# So we need to filter it out
df = df[df['is_interpolated'] == False]

# %% [code]
# check to see if any NaN values are left
df[df[var].isnull()]

# %% [code]
# The most basic curation is to pick a specific time period
df = df.t_filter(start_time = 24, end_time = 48)

# %% [code]
# We can also use interpolate to fill in the missing data points, this can be useful if it's only a few points missing per specimen
df = df.interpolate(variable = 'x', step_size = 60, t_column = 't')

# %% [code]
# We can also group several rows together by increasing the t diff, here we increase
# from 60 to 120, so we find the average of every two rows if there
df = df.bin_time(column = var, bin_secs =  120, function = 'mean')

# %% [code]
# Once we've completed our curation or if we just want to remove specimens that don't have enough values,
# we can call curate to remove all specimens with too few points still
df = df.curate(points = ((24*60*60) / 60)-1) # 24*60*60 = seconds in a day and then divided by are t diff per row. 

# Note: The interpolate method returns rows 1 shorter than before so you'll need to add a minus 1 if using curate after
# Note: If you've called the above bin_time this curate will return an empty dataframe

# %% [code]
# When using x position data the interesting part is how the fly positions itself in relation to the food
# However this will be different on the x,y axis for flies on either side of the ethoscope, so lets normalise it
# You only need to run this is using the x variable
df_r = df.xmv('region_id', list(range(11,21)))
df_l = df.xmv('region_id', list(range(1,11)))
df_r['x'] = 1 - df_r['x']
df = df_l.concat(df_r)

# %% [markdown]
## 3. Normalise the data

# %% [markdown]
#### The ethoscope data can do with a bit of augmentation to make it perform better in HCTSA

# %% [code]
# First lets put our data into numpy arrays
list_x = df.groupby(df.index, sort = False)[var].apply(list)
arr_x = np.array([np.array(x) for x in list_x])
# Here we grab the ids of each for the labels that we'll use later
list_id = list_x.index.tolist()

# %% [code]
# Use some or all of these functions to normalise the data between specimens

# norm transforms the data to be between 0 and 1
def norm01(x):
    return (x-np.nanmin(x))/(np.nanmax(x)-np.nanmin(x))
# or
# find the zscore
def zscore(x):
    return (x-np.mean(x))/(np.std(x))

# Only use this if looking at phi, it changes it be only from 0-90 or horizontal to veritcal as the ethoscope doesn't track direction
def norm_phi(x):
    return np.where(x > 90, 90 - (x - 90), x)

# Smooth out the time series data
def moving_average(a, n) :
    ret = np.cumsum(a, dtype=float)
    ret[n:] = ret[n:] - ret[:-n]
    return ret[n - 1:] / n

# %% [code]
# We can have a look at this effect through some plots
import matplotlib.pyplot as plt

# %% [code]
ind1 = np.random.randint(len(arr_x))
ind2 = np.random.randint(len(arr_x))

plt.figure(figsize=(9, 6))
plt.plot(arr_x[ind1], color = 'red')
plt.plot(arr_x[ind2], color = 'blue')
plt.xlabel('Time (s)', fontsize=12)
plt.ylabel(f'{var} value', fontsize=12)
plt.show()

# %% [code]
# Apply the function along the axis
arr_x = np.apply_along_axis(norm, 1, arr_x)
arr_x = np.apply_along_axis(partial(moving_average, n = 5), 1, arr_x)

# %% [code]
plt.figure(figsize=(9, 6))
plt.plot(arr_x[ind1], color = 'red')
plt.plot(arr_x[ind2], color = 'blue')
plt.xlabel('Time (s)', fontsize=12)
plt.ylabel(f'{var} value', fontsize=12)
plt.show()

# %% [markdown]
## 4. Create the Time Series, Keywords, and labels for the .mat file

# %% [code]
# Here the time series data is augmented to fit into the correct format
# it's stored as the variable ts_mat
ts_mat = np.zeros((len(arr_x),), dtype = object) 
for i in range(len(arr_x)):
    ts_mat[i] = arr_x[i]

# %% [code]
# HCTSA allows keywords to describe each series, we can grab these from the meta data
# Feel free to make your own lists of keywords if you haven't got them in your metadata, just make sure they have the right postions in relation to the ids
# Filter by the list_id to make sure we only grab those we retained from the data curation and it's the correct order
list_sex = df.meta['sex'].filter(list_id).tolist()
list_sle = df.meta['sleep_deprived'].filter(list_id).tolist()

# %% [code]
# Add the keywords together into a single string to match the HCTSA format
list_key = np.array([f'{i},{q}' for i, q in zip(list_sex, list_sle)], dtype=object)

# %% [code]
# Make the id/label list into an object tyoe numpy array so it formats properly
list_lab = np.array(list_id, dtype=object)

# %% [code]
# Save to a .mat file
savemat('./ts.mat', {'timeSeriesData' : ts_mat, 'labels' : list_lab, 'keywords' : list_key})
```
