## Supplementary material for "ethoscopy & ethoscope-lab: a framework for behavioural analysis to lower entrance barrier and aid reproducibility": Jupyter Notebooks: 6_Ethoscopy_catch22_tutorial.ipynb

```
# %% [markdown]
## Tutorial number 6
### Catch22 analysis of ethoscope data using ethoscopy

# %% [markdown]
Catch22 is a shortened version of HCTSA, a time series comparative analysis toolbox in matlab. Catch22 has been adapted to also work in python and uses the top 22 most used analytical tests from the 100s used in HCTSA. 

See heere for more details: https://feature-based-time-series-analys.gitbook.io/catch22-features/
or their github: https://github.com/DynamicsAndNeuralSystems/pycatch22

# %% [markdown]
## 1. Load the dummy dataset

# %% [code]
import ethoscopy as etho
import pandas as pd
import numpy as np
import pycatch22

# %% [code]
# Load in the data from the circadian tutorial
from ethoscopy.misc.get_tutorials import get_tutorial
data, metadata = get_tutorial('circadian')
df = etho.behavpy(data, metadata, check = True)

# %% [markdown]
## 2. Some data curation

# %% [code]
# We'll look at movement here
var = 'moving'

# %% [code]
# The most basic curation is to pick a specific time period
df = df.t_filter(start_time = 0, end_time = 9*24)

# %% [code]
# We can also use interpolate to fill in the missing data points, this can be useful if it's only a few points missing per specimen
df = df.interpolate(variable = var, step_size = 60, t_column = 't')

# %% [code]
# Or we can also group several rows together by increasing the t diff, here we increase
# from 60 to 120, so we find the average of every two rows

## We won't run this here, but keep it in mind for the future if you have too few data points 
# df = df.bin_time(column = var, bin_secs =  120, function = 'mean')

# %% [code]
# Call .summary() to check the amount of data points
df.summary(True)

# %% [code]
# Once we've completed our curation or if we just want to remove specimens that don't have enough values,
# we can call curate to remove all specimens with too few points still
# We can see from summary() that the normal amount is 11999
df = df.curate(points = 11999)

# Note: The interpolate method returns rows 1 shorter than before so you'll need to add a minus 1 if using curate after
# Note: If you've called the above bin_time this curate will return an empty dataframe

# %% [code]
# When using x position data the interesting part is how the fly positions itself in relation to the food
# However this will be different on the x,y axis for flies on either side of the ethoscope, so lets normalise it
# You only need to run this is using the x variable, which we aren't right now
# df_r = df.xmv('region_id', list(range(11,21)))
# df_l = df.xmv('region_id', list(range(1,11)))
# df_r['x'] = 1 - df_r['x']
# df = df_l.concat(df_r)

# %% [markdown]
## 3. Normalise the data

# %% [markdown]
#### The ethoscope data can sometimes do with some augmentaion to make sure it perfroms better in Catch22
#### Try running your data unnormalised and normalised, and with different methods to see the results

# %% [code]
# Catch22 takes only lists, but to normalise the data we'll have to put it into a numpy array first
list_x = df.groupby(df.index, sort = False)[var].apply(list)
arr_x = np.array([np.array(x) for x in list_x])
# Here we grab the ids of each for the labels that we'll use later
list_id = list_x.index.tolist()

# %% [code]
# Use some or all of these functions to normalise the data between specimens

# norm01 transforms the data to be between 0 and 1
def norm01(x):
    return (x-np.nanmin(x))/(np.nanmax(x)-np.nanmin(x))
# or
# find the zscore
def zscore(x):
    return (x-np.mean(x))/(np.std(x))

# Only use this if looking at phi, it changes it be only from 0-90 or horizontal to veritcal as the ethoscope doesn't track direction
def norm_phi(x):
    return np.where(x > 90, 90 - (x - 90), x)

# Smooth out the time series data
def moving_average(a, n) :
    ret = np.cumsum(a, dtype=float)
    ret[n:] = ret[n:] - ret[:-n]
    return ret[n - 1:] / n

# %% [markdown]
#### As the data here is just 1s and 0s (True and False) we don't want to perform any curation. Head to tutorial 6 for information on how to apply it to other sets of data. But for now just bear in mind that it's a useful pre-processing tool.

# %% [markdown]
## 4. Run Catch22 and store in a behavpy dataframe 

# %% [code]
# Here the time series data is augmented to fit into the correct format, a nested list, and run through the catch22 function: pycatch22.catch22_all()
# You can also call individual tests if you know which ones you need, see the pycatch documents for more details
# If you want the mean and std then call catch24 = True within the function 
data = [pycatch22.catch22_all(list_x[i])['values'] for i in range(len(list_x))]
cols = pycatch22.catch22_all(list_x[0])['names']

c22 = pd.DataFrame(data, columns = cols, index = list_id)

# %% [markdown]
#### To compare the scores and find trends we need to normalize and standardise the data

# %% [code]
# sklearn is a common python package for statistics and machine learning, if you dont have in installed, install from pip now.
from sklearn.preprocessing import normalize, MinMaxScaler

# %% [code]
# We call the normalisation across each variable (column)
# normalize() scales vectors to a unit norm so that the vector has a length of 1
normed = normalize(c22, axis=0)

# %% [code]
# We scale the data so all scores are fitted to between 0 and 1 (classification techniques perforn much better with same scaled data)
scaled = MinMaxScaler().fit_transform(normed)

# %% [code]
# Lets put the data into a dataframe and then a behavpy dataframe
c22_norm = pd.DataFrame(scaled, columns = c22.columns, index = list_id)
c22_norm.index.name = 'id'
c22_df = etho.behavpy(c22_norm, df.meta)

# %% [markdown]
## 5. Viewing the results

# %% [code]
# We'll use matplotlib and its add on pacakage seaborn here to create graphs rather than the usual plotly
from matplotlib import pyplot as plt
# You may need to pip install seaborn
import seaborn as sns

# %% [markdown]
#### Heatmaps are often the best way to get a quick view of where your outputs vary

# %% [code]
heatmap = sns.heatmap(c22_df, cmap='PiYG')
heatmap.set_title('Heatmap of normed output', fontdict={'fontsize':12}, pad=12);

# %% [markdown]
#### We can see a the two histogram mode tests all score roughly the same, so they're not likely to be useful. However, both the CO seem to have a variation in scores across the board. Lets look at the heatmap for each group we have in the metadata for circadian length: short, WT, and long

# %% [code]
# We iterate and filter with xmv to get each group, removing the y ticks so its less cluttered 
f, ax = plt.subplots(3,1, sharex='col')
f.subplots_adjust(hspace=0.3)

for c, typ in enumerate(['short', 'wt', 'long']):
    tmp_df = c22_df.xmv('period_group', typ)
    g = sns.heatmap(tmp_df, cmap="PiYG", cbar=False, ax=ax[c])
    g.set(yticklabels=[])  
    g.set(title=typ)
    g.set(ylabel=None)
    g.tick_params(left=False)

# %% [markdown]
#### We can see clearly the difference between the groups with the CO analysis. As well as big differences between long to WT and short when looking at FC local Simple.

# %% [code]
# We can also plot the correaltion bewteen each variable to see the relationship between them all

heatmap = sns.heatmap(c22_df.corr(), vmin=-1, vmax=1, cmap = sns.diverging_palette(220, 10, as_cmap=True))
heatmap.set_title('Correlation Heatmap', fontdict={'fontsize':12}, pad=12);

# %% [markdown]
#### There are strong clusters of tests that are similiar to each other which is to be expected. But we can also see CO ters are strong correlated both postively and negatively with many of the others

# %% [code]
# We'll need the labels to colour code the plot, but to make sure they have the same order we'll match them to the main df and remove them
# The order in the metadata is often (and should be) the same as order in the data, but sometimes with curation it can change so this is just to be safe
# We make a dictionary of the id and the label and map it to data
label_dict = c22_df.meta['period_group'].to_dict()
c22_df['label'] = c22_df.index.to_series().map(label_dict)

# %% [markdown]
#### A pairwise plot is a more visual version of th correlation plot, but we can also colour by our labels and see how they group

# %% [code]
# Pairwise plot will give you the most insight into which variables have distinct populations per label
sns.pairplot(c22_df, vars = cols, hue = 'label')

# %% [markdown]
#### We can see just visually from the pairwise plot what most of the variable combinations serperate the populations into roughly distint groups, which bodes well for classification. We can evene see a few with linear correlations such as IN_AutoMutualInfoStats_40_gaussian_fmmi X CO_FirstMin_ac.

# %% [markdown]
#### From the pairwise plot we can see that the tests can likely seperate the groups well. We can visualise this all together with a PCA or tSNE plots. Both theses plots take all the variables are reduce them to a lower dimmentiality at the cost of some detail, often 2 or 3 as we can then visulise them. In the reduction the algorithm will creat new components that represent the most valuable variables.

#### There are a few differences between PCA and tSNE, but in short tSNE works better with non-linear relationships and can group nearby points better. However, it does this at the cost of true distance between groups and over represent differences between clusters.

# %% [code]
# import both functions from sklearn
from sklearn.manifold import TSNE
from sklearn.decomposition import PCA

# %% [code]
# lets turn each row into a list and nest them, we also need to drop the labels as they are not numerical
X = c22_df.drop(columns = ['label']).values.tolist()
X = np.array([np.array(i) for i in X])

# %% [code]
# We'll intialise the PCA function and state we want it reduced to 2 variables
pca = PCA(n_components=2)
# fit_transform() learns the data and then transforms it to the 2 principle components.
X_pca = pca.fit_transform(X)

# %% [code]
# Turn the output into a pandas datafram and add the labels
pca_df = pd.DataFrame(X_pca, columns = ['PCA1', 'PCA2'])
pca_df['label'] = c22_df['label'].tolist()

# %% [code]
plt.figure(figsize=(10,4))

sns.scatterplot(data=pca_df, 
                x="PCA1", 
                y="PCA2", 
                hue="label")

plt.title("PCA Plot",
        fontsize=14)
plt.xlabel('First Principal Component',
        fontsize=10)
plt.ylabel('Second Principal Component',
        fontsize=10)

# %% [markdown]
#### We can see the PCA analysis has 3 distinct groups with some of the WT labelled flies in the short cluster. All long labelled flies are seperate and easy to distinguish.

# %% [code]
# Repeat what we did above but with the TSNE function
tsne = TSNE(n_components=2, random_state=42)
X_ts = tsne.fit_transform(X)
ts_df = pd.DataFrame(X_ts, columns = ['TS1', 'TS2'])
ts_df['label'] = c22_df['label'].tolist()

# %% [code]
plt.figure(figsize=(10,4))
 
sns.scatterplot(data=ts_df, 
                x="TS1", 
                y="TS2", 
                hue="label")
 
plt.title("TSNE Plot",
        fontsize=14)
plt.xlabel('First TSNE component',
        fontsize=10)
plt.ylabel('Second TSNE component',
        fontsize=10)

# %% [markdown]
#### A similar output to the PCA with long being clustered seperately and some mixing of the short and WT labels in two clusters.

# %% [markdown]
## 6. Classification

# %% [markdown]
#### From the above plots it looks like the dataset is good for some classification attempts, we'll go through two of the powerful but simple techniques SVM and Random Forest classifier.

# %% [code]
# We need to load in a few more bits from sklearn
from sklearn.svm import SVC
from sklearn.model_selection import train_test_split
from sklearn.pipeline import Pipeline
from sklearn.model_selection import GridSearchCV
from sklearn.metrics import accuracy_score, confusion_matrix, classification_report
from sklearn.preprocessing import LabelEncoder
from sklearn.ensemble import RandomForestClassifier

# %% [code]
# Lets put our labels into variable y
y = c22_df['label'].tolist()
# And split our dataset into both a training set (70%) and testing set (30%)
# Stratify makes sure there is the same ratio of all groups in both train and test
X_train, X_test, y_train, y_test = train_test_split(X, y, test_size=0.3, random_state=12, stratify = y)

# %% [code]
# This function prints out accuracy and reports of the classifier
def print_score(clf, X, y):
        pred = clf.predict(X)
        clf_report = pd.DataFrame(classification_report(y, pred, output_dict=True))
        print(f"Accuracy Score: {accuracy_score(y, pred) * 100:.2f}%")
        print("_______________________________________________")
        print("Classification Report:")
        print(clf_report)
        print("_______________________________________________")
        print("Confusion matrix:")
        print(confusion_matrix(y, pred))

# %% [markdown]
#### Support Vector Machines (SVM) is a machine learning algorithm that creates a decision boundry in the variable space that it thinks best seperates your given groups. We'll kick off by just calling the most basic SVM, one that employs a linear kernel.

# %% [code]
# Create the classifier and fit it to the data
svm_clf = SVC(kernel='linear')
svm_clf.fit(X_train, y_train)
# The train dataset score
print_score(svm_clf, X_train, y_train)

# %% [code]
# The test dataset score
print_score(svm_clf, X_test, y_test)

# %% [markdown]
#### The linear SVM performs very well on the training dataset (which is to be assumed), but has a poor score for the test dataset. We could play around with the settings for SVM function manually or we can perform a grid search. A grid search takes a list of different inputs to the classifying function and runs all the combination, finding the one with the best score.

# %% [code]
# First create a dictionary of the parameters you want to change and the list of variabels to apply
# We'll use all the kernels avaiable to us and a range of gamma and C
# Head to the sklearn wiki for information on tthe parameters
param_grid = {'C': [0.01, 0.1, 0.5, 1, 5, 10, 100], 
            'gamma': [1, 0.75, 0.5, 0.25, 0.1, 0.01, 0.001, 'auto'], 
            'kernel': ['rbf', 'poly', 'linear']} 
# Now we setup the Grid search with the SVM, the parameters. CV is a resampling method that test and trains within the given dataset
grid_svm = GridSearchCV(SVC(), param_grid, verbose=True, cv=3)

# %% [code]
# Fit the data to the gridsearch
grid_svm.fit(X_train, y_train)
params_svm = grid_svm.best_params_
print(f"Best params: {params_svm}")

# %% [code]
# Lets take the parameters and score then
svm_clf = SVC(**params_svm)
svm_clf.fit(X_train, y_train)
print('Training score:')
print_score(svm_clf, X_train, y_train)
print('Testing score:')
print_score(svm_clf, X_test, y_test)

# %% [markdown]
#### The grid searched parameters perform worse on the trained dataset, but better on the test set. Meaning the it's not as overfitted to the training set!

# %% [markdown]
#### Lets have a look at an SVM when applied to the PCA results from above to gain a visual understanding of what's going on

# %% [code]
# We need to encode the labels as numbers to colour code them with the plotting function below
le = LabelEncoder()
le.fit(['short', 'wt', 'long'])
y2 = le.transform(pca_df.label.to_list())

# %% [code]
# We'll take the values for the PCA dataframe, and the set the labels a yp
Xp = pca_df.values[:, :2] 
yp = y2

# Creates a grid for the plot points to applied to
def make_meshgrid(x, y, h=.02):
    x_min, x_max = x.min() - 0.1, x.max() +0.1
    y_min, y_max = y.min() - 0.1, y.max() + 0.1
    xx, yy = np.meshgrid(np.arange(x_min, x_max, h), np.arange(y_min, y_max, h))
    return xx, yy

# This function plots the different areas the SVM decides for each label in the variable space
def plot_contours(ax, clf, xx, yy, **params):
    Z = clf.predict(np.c_[xx.ravel(), yy.ravel()])
    Z = Z.reshape(xx.shape)
    out = ax.contourf(xx, yy, Z, **params)
    return out

# Call a grid search on this PCA values and call the best fit model
grid_svm.fit(Xp, yp)
best_params = grid_svm.best_params_
model = SVC(**best_params)
clf = model.fit(Xp, yp)

fig, ax = plt.subplots()
title = ('Decision surface of the best fit SVM')
# Set-up grid for plotting.
X0, X1 = Xp[:, 0], Xp[:, 1]
xx, yy = make_meshgrid(X0, X1)

# Plot the svm areas and variable points
plot_contours(ax, clf, xx, yy, cmap = 'coolwarm', alpha=0.8)
ax.scatter(X0, X1, c=yp, s=20, cmap = 'coolwarm', edgecolors='k')
ax.set_ylabel('PCA 2')
ax.set_xlabel('PCA 1')
ax.set_xticks(())
ax.set_yticks(())
ax.set_title(title)
plt.show()

fig.savefig("./figure3c.svg")

# %% [markdown]
#### This visuliases nicely the problem the SVM is having with classifing at 100%, it's the points we noticed above that are WT but have short sleeping like phenotype

# %% [markdown]
#### Next we'll look at random forest classifers (RFC). RFC's are a collection of tree classifiers, each tree classifier works on a subset of the data, making decisions at each leaf to bag the data into one of the labels. On their own they can often overfit to the training dataset, however as a forest and added randomness this can be accounted for. Random forests often perform the best for a wide range of datasets.

# %% [code]
# Lets intialise the classifer and create a grid search for the off
# Head to the sklearn wiki to understand the parameters
rfc = RandomForestClassifier(random_state=42)
param_grid = {'n_estimators': [100, 200, 300, 500],
            'max_features': ['sqrt', 'log2'],
            'max_depth' : [4,5,6,7,8,10],
            'criterion' :['gini', 'entropy']}
grid_rfc = GridSearchCV(rfc, param_grid, refit=True, verbose=1, cv=3)

# %% [code]
# Call and fit the grid search for the RFC
grid_rfc.fit(X_train, y_train)
params_rfc = grid_rfc.best_params_
print(f"Best params (RFC): {params_rfc}")

# %% [code]
# Lets see how it's done
rfc_clf = RandomForestClassifier(**params_rfc, random_state = 42)
rfc_clf.fit(X_train, y_train)
print('Scores for the trained dataset:')
print_score(rfc_clf, X_train, y_train)
print('Scores for the test dataset:')
print_score(rfc_clf, X_test, y_test)

# %% [markdown]
#### Sadly it's not performed as well as the best SVM. We could play around with more parameters, but for now it looks like the SVM score of 80% is the best we'll get (which is still pretty good).

# %% [markdown]
#### If you'd like to perform more tests look online for tutorials about the other classifiers in the sklearn ecosystem. You can also head to the HCTSA tutorial page and look to copy some of their techniques from matlab to python: https://hctsa-users.gitbook.io/hctsa-manual/
```
