## Supplementary material for "ethoscopy & ethoscope-lab: a framework for behavioural analysis to lower entrance barrier and aid reproducibility": Jupyter Notebooks: 7_Working_with_custom_data.ipynb

```
# %% [markdown]
## Tutorial number 7
### Working with custom (non ethoscope) dataset

This tutorial shows how to import data that are not coming from ethoscopes or Drosophila activity monitors.
We will load a CSV file collecting human FitBit activity data from 35 individuals over a month (1 point per day)

# %% [code]
import ethoscopy as etho

import pandas as pd
import numpy as np
np.random.seed(4843094)

# This tutorial requires version 1.1.7 or greater
etho.__version__

# %% [code]
# The fitbit dataset was downloaded from https://www.kaggle.com/datasets/singhakash/fitbit-dataset?resource=download 
# The file contains 32 days of human activity such as calories, steps, distance in meters, floors, 
# minutes sitting, minutes of moderate activity, minutes of intense activity as well as the calories burned for the activities
# for 35 individuals of which we have no information. The data was gathered with a Fitbit Charge HT fitness tracker and every observation regards one day.
fitbit_dataset = "./FitBit_data.csv"

# First we need to read the csv file into a pandas.DataFrame
data = pd.read_csv(fitbit_dataset)

# Then we specify which column should be used as index
data = data.rename(columns={"Id": "id"}).set_index("id")

# These sample fitbit data come with no metadata so we generate some random metadata just as proof of principle
metadata = pd.DataFrame({
                            "id" : data.index.unique(),
                            "age" : np.random.randint(20, 30, size=data.index.unique().size),
                            "sex" : np.random.choice(["Male", "Female"], size=data.index.unique().size)
                        }).set_index("id")

# %% [code]
# Now we load both into an ethoscopy behavpy object using the seaborn canvas
df = etho.behavpy(data, metadata, check = True, canvas = 'seaborn')
df.head()

# %% [code]
# And we can start working with it. Keep in mind these is just a proof of principle. 
# Lots of methods will not be working because ethoscopy expects more than just one point of data per day
fig, stats = df.plot_quantify("VeryActiveMinutes", facet_col="sex")

# %% [code]
```
