## Supplementary material for "ethoscopy & ethoscope-lab: a framework for behavioural analysis to lower entrance barrier and aid reproducibility": Jupyter Notebooks: notebook_paper.ipynb

```
# %% [markdown]
# ethoscopy & ethoscope-lab: a framework for behavioural analysis to lower entrance barrier and aid reproducibility

# %% [markdown]
## Figure 1

# %% [markdown]
## 1. Loading the data

# %% [code]
import ethoscopy as etho
import pandas as pd

# %% [code]
# This tutorial required version 1.1.8 or greater
etho.__version__

# %% [code]
# import this function to get the tutorial dataset
from ethoscopy.misc.get_tutorials import get_tutorial

# %% [code]
# We'll be using the same dataset as in the overview tutorial, so use the same function below with the argument 'overview' 
# Load the data and metadata, and then intialise it into a behavpy_HMM 
data, metadata = get_tutorial('overview')
df = etho.behavpy_HMM(data, metadata, check = True)

# %% [code]
df = df.t_filter(start_time = 24, end_time = 144)

# %% [markdown]
### (a)

# %% [code]
fig = df.xmv('sleep_deprived', False).heatmap(variable = 'asleep', title = 'Control')
fig.show()

# %% [code]
# As we saw in the overview tutorial the dataset we have has half the specimens sleep deprived later in the experiment
fig = df.xmv('sleep_deprived', True).heatmap(variable = 'asleep', title = 'Sleep Deprived')
fig.show()

# %% [markdown]
### (b)

# %% [code]
fig = df.t_filter(start_time = 24, end_time = 96).plot_overtime(variable = 'asleep', wrapped = True, title = 'Baseline Sleep')
fig.show()

# %% [markdown]
### (c)

# %% [code]
# Ethoscopy also has pretrained HMMs for both Male and Female CS Drosophila, trained on several days worth of data on hundreds of flies
from ethoscopy.misc.get_HMM import get_HMM

# Have the argument as "M" or "F" for the male or female dataset respectively
h = get_HMM("M")

# %% [code]
fig = df.plot_hmm_overtime(
            hmm = h, 
            variable = 'moving', 
            wrapped = True, 
            labels = ['Deep sleep', 'Light sleep', 'Quiet awake', 'Full awake'], 
            bin = 60, 
            title = 'Hidden Markov Analysis of sleep stages'
            )
fig.show()
```
